# A characterization of mouse retinal ganglion cell types labeled with AAV tools

**DOI:** 10.1101/2025.06.02.657062

**Authors:** Seoyoung Son, Deborah Langrill Beaudoin, Abdul Rhman Hassan, Magaret Sena Akpo, Tomomi Ichinose, Andrew M Garrett

## Abstract

The mouse retina is made-up of approximately 150 types of neurons each with unique characteristics and functions in interpreting visual information. Recent efforts to categorize cell types using molecular markers, morphology, and electrophysiological response properties have provided a wealth of information and a host of tools for studying specific cell types. AAV-based approaches have several advantages over transgenic mouse lines, including ease of application to many different animal models without extensive crossing and their amenability to intersectional approaches. Here, we provide an in-depth characterization of retinal ganglion cell types labeled by two AAV vectors drawn from a recent panel of constructs with synthetic promoters. Each promoter analyzed here was derived from a gene expressed in a cell type specific manner. Using a combination of morphology, molecular markers, and electrophysiological measurement of light responses, we found that each vector labeled distinct subsets of RGCs. However, both labeled more cell types than expected from the expression pattern of the promoter’s endogenous gene. We then characterized the projection patterns of these RGC types to the brain, finding that each AAV type labeled distinct axonal populations. These tools provide new access to a unique subset of cells and will be instrumental to future studies analyzing their functions and connectivity.

## Introduction

The vertebrate retina is a highly organized neural tissue responsible for the initial stages of visual processing (Demb and Singer 2015; Gollisch and Meister 2010). The mouse retina closely mirrors the human retina in both structure and function, making it an invaluable model for studying visual circuitry and disease (Hahn et al. 2023). The mouse retina is a laminated neural tissue composed of five principal neuronal cell types: photoreceptors, horizontal cells (HCs), bipolar cells (BCs), amacrine cells (ACs), and retinal ganglion cells (RGCs). These cell types are distributed across distinct layers, with photoreceptor cell bodies located in the outer nuclear layer (ONL), interneurons (horizontal, bipolar, and amacrine cells) residing in the inner nuclear layer (INL), and RGCs positioned in the ganglion cell layer (GCL). RGCs, the principal output neurons of the mouse retina, receive and integrate visual input from photoreceptors through interneurons, subsequently projecting axons to various regions of the brain including the lateral geniculate nucleus (LGN), sensory related superior colliculus (SCs), suprachiasmatic nucleus (SCH), and more than 40 additional brain areas throughout the brain (Beier et al. 2021). Characterizing RGC types is crucial for understanding visual processing, mapping neural circuits, ultimately aiding in more effective disease research and therapeutic strategies.

In mice, approximately 45 distinct RGC subtypes have been identified based on morphological, molecular, and functional features. Three major public databases have contributed significantly to this classification: EyeWire Cell Museum (http://museum.eyewire.org) built using serial electron microscopy and crowd-sourced dendritic reconstructions (Bae et al. 2018). Mouse Retina Cell Atlas (MRCA), based on single-cell RNA sequencing, clustered RGCs by gene expression profiles (Li et al. 2024). RGCtype.org, which integrates transcriptomic, anatomical, and electrophysiological data to define functionally relevant RGC subtypes (Goetz et al. 2022).

While transgenic mouse models have enabled targeting of specific RGC populations, for example, *Jam2-creERT2* for J-RGCs (Kim et al. 2008), *Cart-TG1-Cre* (Martersteck et al. 2017) and *Pcdh9-Cre* (Lilley et al. 2019) for direction-selective ganglion cells (DSGCs), and *Etv1-creERT2*, *Kcng4-Cre* (Krieger et al. 2017), or *Thy1-YFP (W3)* (Zhang et al. 2012) for alpha and W3-RGCs, these models are often limited in their translational relevance due to the requirement for genetic modification. Further, using them experimentally is limited by the need to maintain and cross multiple lines if one wants to analyze multiple cell types. In contrast, adeno-associated viruses (AAVs) offer a versatile, efficient, and species-translatable alternative for targeting specific cell types. AAVs are widely used in both basic research and clinical settings due to their safety profile, ease of production, and ability to deliver genetic material across species. Cell-type-specific targeting in the retina using AAVs has been achieved by placing cell-selective promoter sequences upstream of reporter or effector genes (Jüttner et al. 2019). However, the systematic evaluation of RGC subtype specificity for such AAV-delivered promoters has remained limited.

In this study, we investigated two synthetic AAV promoters, ProA13 (derived from *Nppb*) and ProA27 (from *Serpinb1b*), each driving membrane-targeted GFP expression. We evaluated the RGC subtypes labeled by these promoters through a through an integrated approach combining molecular, morphological, and physiological analyses. First, we performed immunohistochemistry to identify molecular markers of specific RGC subtypes. Next, we conducted morphological analysis using ChAT band-referenced dendritic reconstructions. Then, we performed patch-clamp electrophysiology to correlate functional responses with individual RGC morphologies. Finally, we examined central projection patterns of ProA13- and ProA27-labeled RGCs to characterize their downstream targets. These findings demonstrate that AAVs equipped with distinct synthetic promoters can selectively target defined subsets of retinal ganglion cell (RGC) subtypes. While each promoter drives expression in a limited number of RGC types, this approach offers a practical and flexible tool for cell type–specific manipulation and for studying both image-forming and non-image-forming visual circuits.

## Results

### ProA13 drive expression of ganglion cells, amacrine cells, and horizontal cells while ProA27 mainly target ganglions cells

Jüttner et.al. created a library of 230 AAV vectors with different strategies to achieve cell type specificity. Their A-type strategy was to use predicted promoter sequence from genes with cell-type-specific expression (Jüttner et al. 2019). We chose two vectors from this class that they reported to express 1) only in RGCs and 2) in a sparse enough population that they could label one or a few specific types.

Specifically, ProA13 was derived from the *Nppb* gene locus and ProA27 from *Serpinb1b*. Analysis of published single-cell transcriptomic data from the MRCA database, which classifies RGCs into 45 distinct clusters based on gene expression, revealed that *Nppb* is enriched in select intrinsically photosensitive RGCs (ipRGCs) most strongly in M1 (C33/M1 and C40/M1dup) and C31/M2. There was also measurable expression in C22/M5 and ON-type cells in clusters C43/AlphaONS and C38/F-midiON. In contrast, *Serpinb1b* was predominantly expressed in the novel RGC type C11, with additional expression in W3 subtypes (C2/W3D1.2 and C6/W3B), and potential low expression in some ipRGCs (C32, C33, C40) (Fig. 1A).

**Figure 1.**
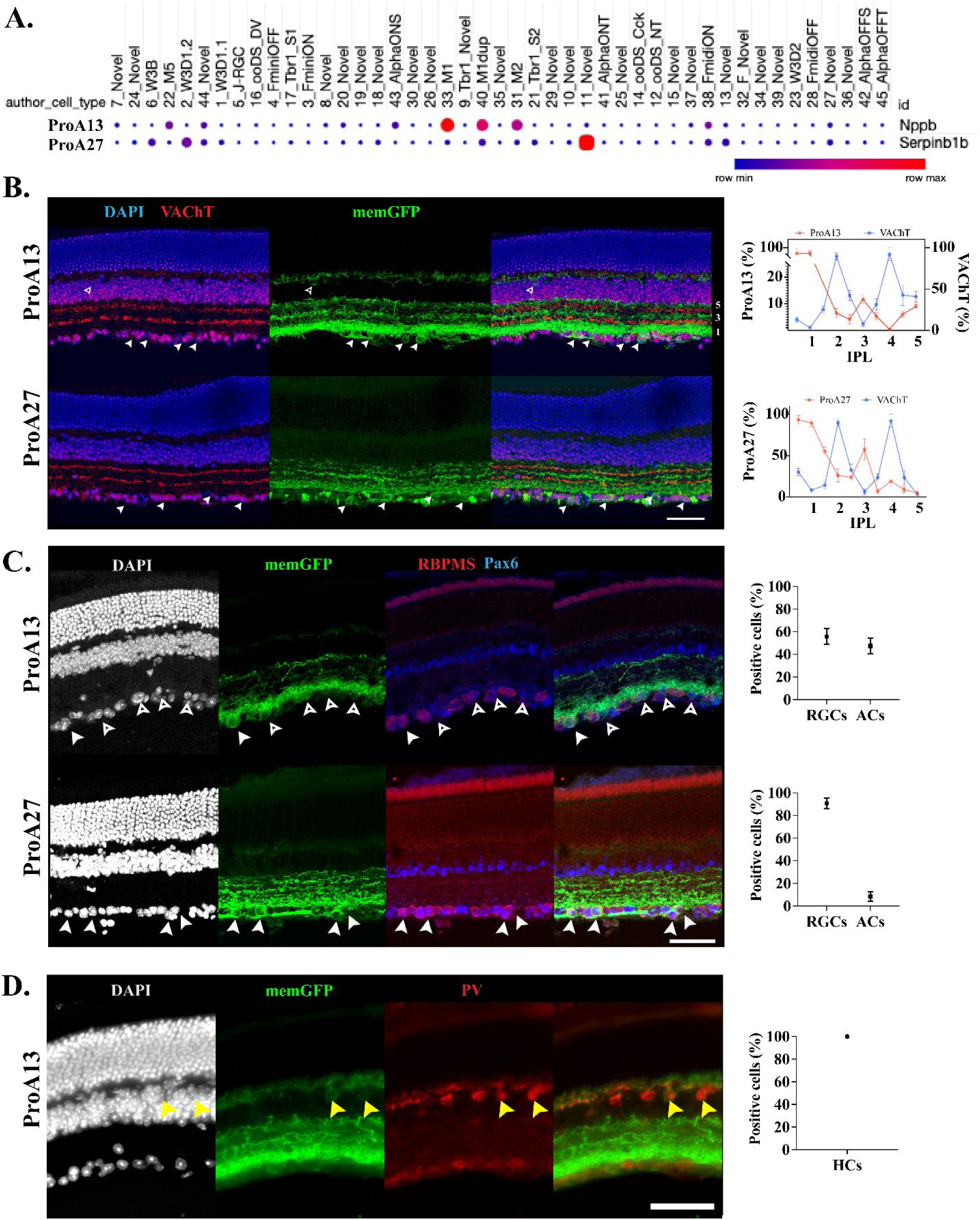
Cell-type specificity of ProA13 and ProA27 AAVs in the mouse retina. (A) Dot plot showing gene expression of *Nppb,* the basis for promoter ProA13 and *Serpinb1b* the basis for ProA27 across 45 transcriptionally defined RGC clusters from the MRCA single-cell RNA-seq dataset. *Nppb* is enriched in ipRGCs (e.g., C22, C31, C33, C40) and ON-RGCs (e.g., C38, C43), while *Serpinb1b* is enriched in W3-RGCs (C2, C6), novel types (C11), and some ipRGCs (C32, C33, C40). Expression intensity is color coded. (B) Representative retinal sections from ProA13 and ProA27 AAV-injected eyes, immunostained for VAChT (Red) to mark cholinergic bands, memGFP (green, white arrowheads), and DAPI (blue). ProA13-RGCs stratify in sublaminae 1, 3, and 5 of the IPL and in the outer margin of the INL (open arrowhead). Stratification patterns in IPL were drawn using IPlaminator ImageJ plugin (Li et al. 2016). ProA13-RGCs were alternating with ChAT bands, while ProA27-RGCs show less distinct stratification with strong signal in layer 3. (C) Immunohistochemistry for RBPMS (red, RGC marker) and Pax6 (blue, displaced amacrine cell marker) in ProA13 and ProA27-AAV-injected retinas. MemGFP-labeled cells in the GCL include both RGCs (PBPMS+ or PBPMS+Pax6+; arrowheads) and ACs (Pax6+; open arrowheads) in ProA13 retinas, but primarily RGCs in ProA27 retinas. Right: Quantification of memGFP+ cells (n = 17-30 images from 3-6 retinas per group, mean ± SEM). (D) Immunolabeling of ProA13-AAVs injected retinas for parvalbumin (PV, red, HC marker) with DAPI (gray) and memGFP (green). memGFP+ cells at the outer INL (yellow arrowheads) colocalize with PV, indicating horizontal cell expression. Right: Quantification of horizontal cell labeling among memGFP+ cells (n = 4 images from 3 retinas, mean ± SEM). Scale bars: 50 μm (B–D).

We subcloned the synthetic promoter sequences upstream of a memGFP reporter coding sequence and packaged the vector in AAV2/2 viral particles. To characterize the *in vivo* targeting profile of each construct, we injected viral particles into the vitreous of adult mice and analyzed memGFP expression in retinal sections. ProA13-labeled cells (ProA13-RGCs) were predominantly located in the GCL and showed dendritic stratification in sublaminae 1, 3, and 5 of the inner plexiform layer (IPL), alternatively positioned with VAChT-positive strata (used as anatomical landmarks at strata 2 and 4). Notably, the sublamina 1 band was broader and brighter than others (Fig. 1B). In contrast, ProA27-labeled cells (ProA27-RGCs) were confined to the GCL and exhibited signal throughout the IPL. However, layer 1 has displayed stronger signal, and a subset of dendrites with bright labeling was observed stratifying in layer To confirm the identity of GFP-positive cells, we performed immunohistochemistry for RBPMS, a pan-RGC marker, and Pax6, which label both displaced ACs and RGCs in the GCL. In ProA13 infected retinas, approximately 52.5% of GFP+ cells in the GCL were Pax6+/RBPMS-, indicating amacrine cell targeting, while the remainder were RGCs (RBPMS+) (Fig. 1C). In ProA27 infected retinas, 91.5% of GFP+ cells in the GCL were RBPMS+, confirming strong RGC specificity. Additionally, we observed ProA13-mediated labeling of cells in the outer margin of the INL, where HCs reside. These cells were identified as parvalbumin (PV)-positive, consistent with HCs (Fig. 1D). No such labeling was observed with ProA27. Together, these findings indicate that ProA13 drives expression in a broader population, including RGCs, displaced ACs, and HCs, while ProA27 exhibits high specificity for RGCs.

### Broader ipRGC Diversity Labeled by ProA27 Than ProA13

Categorizing RGC subtypes is essential for understanding the functional diversity of visual processing pathways (Sanes and Masland 2015). To validate the molecular identity of RGCs labeled by ProA13 and ProA27, we performed immunohistochemistry using antibodies against well-characterized subtype-specific markers: SMI32 for alpha RGCs, melanopsin for ipRGCs, Tusc5 for T5-RGCs, Foxp2 for F-RGCs, and CART for ON-OFF DSGCs (ooDSGCs) (Gallego-Ortega et al. 2022; Hattar et al. 2002; Kay et al. 2011; Rousso et al. 2016; Tran et al. 2019). Among ProA13-labeled cells, approximately 6.1% were positive for melanopsin, while ProA27-labeled cells exhibited a significantly higher melanopsin positivity rate of 14.8% (Fig. 2A). To further resolve the identity of these melanopsin-expressing cells, we performed single-cell morphological reconstructions of melanopsin + RGCs. ProA13-labeled ipRGCs were predominantly M1-type cells, as indicated by their dendritic arbors stratifying exclusively in the outer (OFF) sublamina of the inner plexiform layer (IPL) (Fig. 2B). In contrast, ProA27-labeled melanopsin + RGCs displayed greater subtype diversity, encompassing M1, M2 (ON-stratified), and M3 (bistratified ON-OFF) ipRGCs. Notably, some ON-stratified ipRGCs in the ProA27 population co-expressed SMI32, suggesting their classification as M4 ipRGCs, a subtype known to share molecular features with alpha RGCs (Fig. 2C) (Schmidt et al. 2014). These results demonstrate that ProA27 drives expression in a broader spectrum of ipRGC subtypes than ProA13, enabling access to ipRGC populations with distinct dendritic architectures and potentially diverse functional roles in both image-forming and non-image-forming pathways.

**Figure 2.**
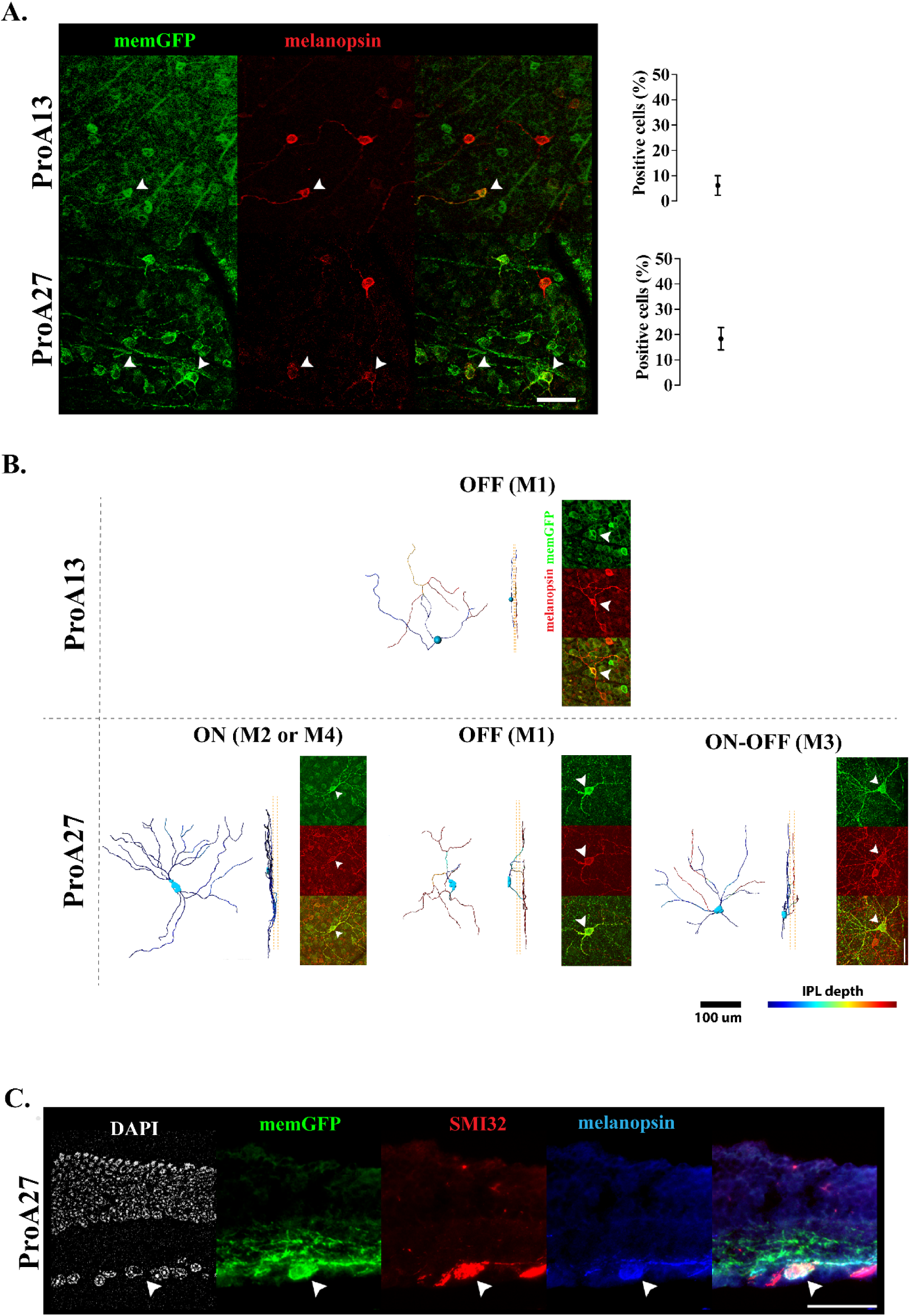
ProA13 and ProA27 label distinct ipRGC subtypes with differing morphological and molecular profiles. (A) Representative immunofluorescence images showing co-localization of memGFP (green) with melanopsin (red) in ProA13- and ProA27-AAVs injected retinas. Arrowheads indicate memGFP+ RGCs that are also melanopsin. Quantification of melanopsin-positive cells among labeled RGCs is shown to the right; ProA27 exhibits a higher proportion of melanopsin + RGCs compared to ProA13. (B) Single-cell reconstructions of melanopsin+ RGCs from ProA13- and ProA27-AAVs injected retinas reveal subtype-specific dendritic stratification in the IPL. ProA13-RGC melanopsin + cell show typical M1 morphology with dendrites confined to the OFF sublamina. In contrast, ProA27-RGCs include M1 (OFF), M2 or M4 (ON), and M3 (bistratified) ipRGC subtypes. Corresponding confocal images confirm subtype identity (melanopsin+) and stratification (yellow dotted line drown at submarine layer 2 and 4 with ChAT signal). IPL depth color coding is shown at bottom right and scale bars was 100 μm. (C) Confocal cross-section from ProA27-injected retina showing colocalization of memGFP (green) with both melanopsin (blue) and SMI32 (red). Arrowheads mark cells co-expressing melanopsin and SMI32, consistent with M4 ipRGC identity. Scale bars for confocal images: 50 μm (A-C).

### ProA13 and ProA27 Label Diverse RGC Subtypes Including Molecularly Defined and Unlabeled Types

Among ProA13-labeled cells, approximately 2.2%, 1.0%, 1.5%, and 0.4% were positive for SMI32, CART, Tusc5, and Foxp2, respectively. In contrast, ProA27-labeled cells exhibited broader subtype coverage, with 14.6% SMI32+, 8.8% CART+, 20.1% Tusc5+, and 2.1% Foxp2+ cells (Fig. 3A–C). ProA27-labeled RGCs displayed well-defined dendritic stratification, particularly within IPL layer 3 (Fig1B). A subset of these layer 3 stratified cells was Tusc5-positive, consistent with the identity of T5-RGCs. Tusc5-negative cells were also present in this layer, indicating molecular heterogeneity within the same stratification domain. Moreover, a subset of ProA27-labeled RGCs co-expressed both Tusc5 and Foxp2, a combination characteristic of F-mini-ON and F-mini-OFF RGCs (Tran et al. 2019). Despite the identification of several well-characterized RGC subtypes, a significant proportion of memGFP-positive cells did not express any of the tested markers. This implies that both ProA13 and ProA27 also label less-characterized or novel RGC types, including Tusc5-negative cells stratified in layer 3 and additional unidentified populations (Fig. 3E). Together, these results highlight the utility of synthetic AAV promoters for accessing a diverse array of molecularly distinct RGCs, including those not readily defined by current molecular identities of RGCs.

**Figure 3.**
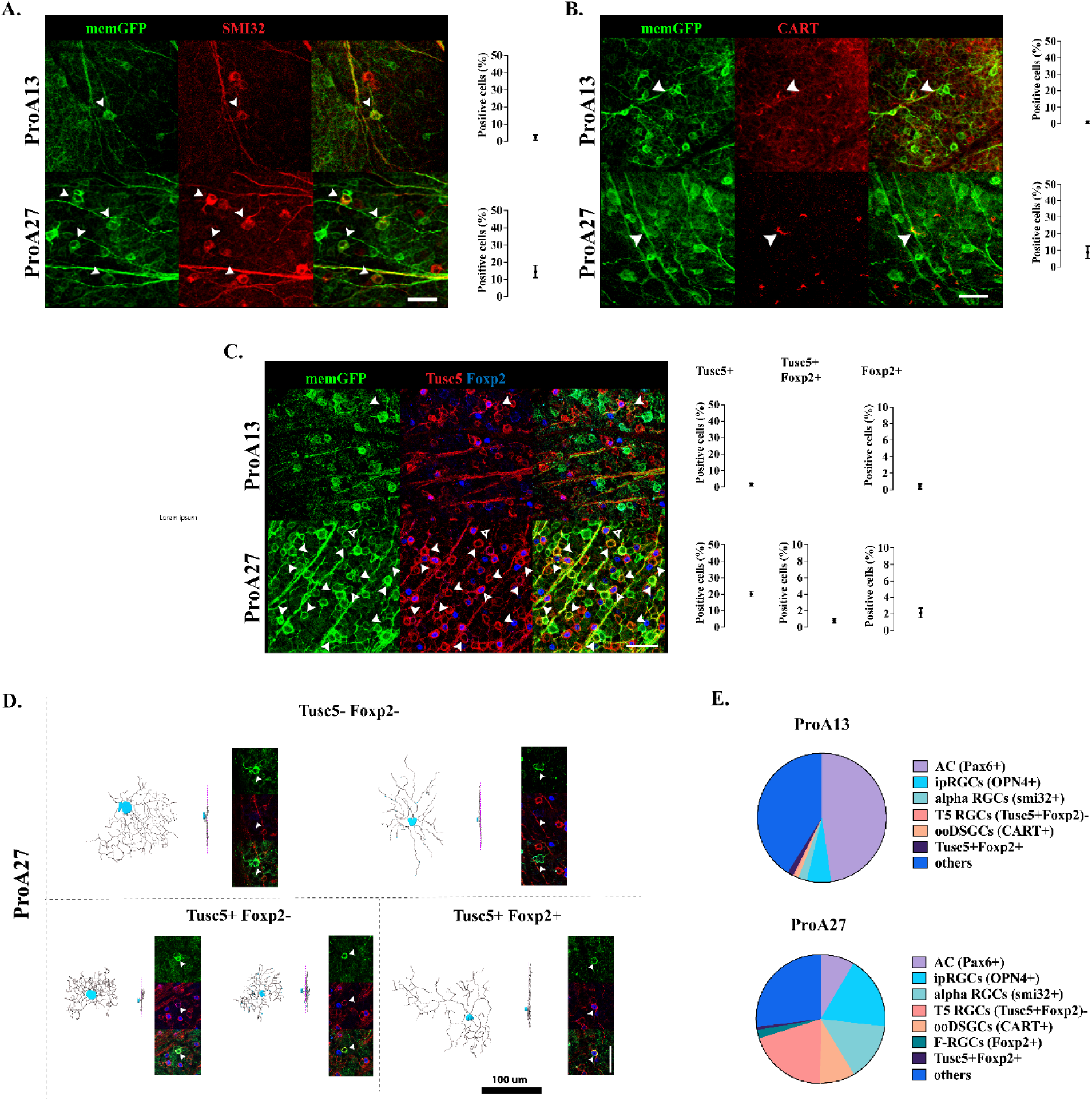
Molecular marker expression and dendritic stratification reveal subtype diversity in ProA13- and ProA27-RGCs. Representative confocal images showing immunostaining of ProA13- and ProA27-RGCs with subtype-specific molecular markers. MemGFP (green) was co-labeled with antibodies against (A) SMI32 (alpha-RGCs; red), (B) CART (ON-OFF direction-selective RGCs; red), and (C) Tusc5 (T5-RGCs; red) and Foxp2 (F-RGCs; blue). Arrowheads indicate double-positive cells (green and red in A-C). Triple-positive cells were indicated with open arrowheads in (C). Quantification of each marker-positive RGCs in ProA13- and ProA27-RGCs are shown to the right. ProA27-RGCs are a greater proportion of each marker than ProA13. (D) Single-cell reconstructions of RGCs stratified in layer 3 by ProA27. Corresponding confocal images confirm subtype identity (tusc5 or Foxp2) and stratification (magenta dotted line drown at submarine layer 3 where tusc5 positive cells dendritic stratification start. (E) Summary of molecular marker coverage among memGFP-positive RGCs. Pie chart illustrating the proportion of ProA13-(left) and ProA27-RGCs (right) positive for each molecular marker. Population in others are suggesting ProA13 and ProA27 also label novel or less well-defined subtypes. Scale bars for confocal images: 50 μm (A-D).

### Morphological and Functional Characterization of RGC Subtypes

To further investigate the subtype identity of RGCs targeted by ProA13 and ProA27, we performed a combined analysis of neuronal morphology and physiological properties. Individual RGCs were reconstructed based on their dendritic arborization and laminar stratification, using ChAT bands as anatomical landmarks to define IPL sublaminae. Morphological parameters, including soma size, dendritic arbor area, and stratification depth, were compared with reference datasets from the EyeWire museum, allowing subtype inference based on established classification schemes. To complement the structural analysis, we conducted patch-clamp electrophysiological recordings, enabling correlation between morphological features and functional light response profiles.

Morphologically, A13-RGCs included various types of ON-type RGCs. One frequently observed type had an arbor area of approximately 20,000 μm stratified adjacent to the RGL. Based on this information and the relative density of dendrite arborization, we linked these to the 9n subtype from the EyeWire dataset. These have been matched to the PixON subtype of ON sustained RGCs (Johnson, Zhao, and Kerschensteiner 2018 and rgctypes.org). Consistent with this, we found multiple cells (n=5) characterized by a sustained spike response to light increments (ON stimuli) and decreased spiking during light decrements (OFF stimuli) (Fig. 4A). Some cells in this group also had a high spontaneous spike activity. Additionally, we measured the spatial tuning using the various spot sizes. The spike rate increased from a small to a larger spot diameter, which peaked at a diameter of 800 µm. Surround inhibition slightly reduced the ON response. No direction selectivity was observed in the recorded cells Another major type stratifying adjacent to the RGL is the ON alpha RGC subtype, which we also found among the ProA13-RGCs. These have a much larger arbor size than PixON and can be identified by their large somas and immunoreactivity for SMI32 (Fig.3). By patch clamp recording we identified cells (n=2) characterized by a transient spiking response at stimulus onset that died away before the end of the stimulus (Fig. 4Biv). Larger spot sizes generally elicited much higher spiking rates, with inhibition beginning to dampen the response at 1000 µm indicating a large receptive field. No direction selectivity was observed in the recorded cells.

**Figure 4.**
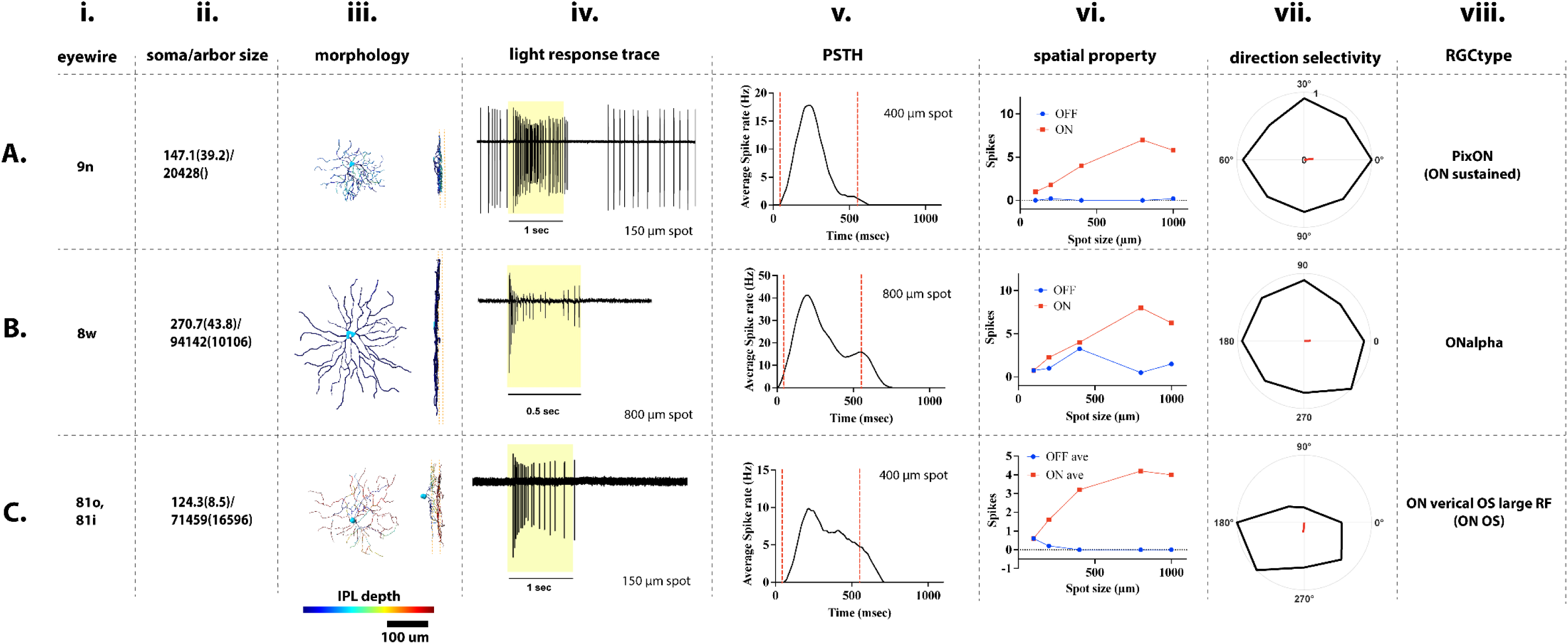
ProA13 labels ON RGC subtypes. (Column i.) Putative cell types were identified based on morphology matching to the EyeWire database. (ii.) Soma and dendritic arbor areas were measured in projections from confocal stacks. (iii.) Examples of traced RGC’s matched to putative EyeWire types shown *en face* and from the side illustrate morphology and dendrite stratification. Patch clamp recordings from a separate set of retinas revealed light response properties including (iv.) spike traces, (v.) average spike rate, or the mean peristimulus time histogram (PSTH), to 1 Hz square-wave modulation with the indicated spot size, (vi.) average total spike counts to 1 Hz square-wave modulation during ON (red) and OFF (blue) plotted as a function of stimulus diameter, and (vii.) vector plot (black) showing total spike count for 8 different directions to a 1 Hz drifting grating with 200 µm bars. Small vector (red) indicated vector sum. These were used to match to (viii.) types in the RGCtype database. (A) Cells morphologically matched to the 9n subtype had an arbor area of ∼20,000 μm^2^ and stratified adjacent to the RGL. Light responses, including raw spike traces (3 cells responding to 1 second 150 µm spot stimulus), spot size preference, and temporal responses were used to match with the PixON subtype. (B) neurons morphologically matched to the 8w type had much larger soma and arbor sizes. Light responses included raw traces responding to a large 800 μm stimulus, spot size preference, and temporal responses suggested on alpha sustained RGC type. (C) a subset of bistratified RGCs more morphologically similar to EyeWire type 81o or 81i. These could correspond to subtype found by patch clamp recording with an ON sustained response pattern. A slight DS preference was seen in this cell.

We also identified cells with bistratified dendrites that corresponded to ON sustained bistratified cells (n=2) characterized by a slightly delayed but fast and sustained spiking rate (Fig. 4C). ON responses increased as spot sizes increased, with a slight decrease after 800 µm indicating a large receptive field. A direction selectivity was observed in response to a full contrast 1 Hz grating with 200 µm bars in 8 directions. Based on morphology and receptive field size, we tentatively assigned these to the ON vertical OS large RF subtype, which corresponds to EyeWire type 81o or 81i (Fig. 4C). Our stimulus did not directly test orientation selectivity in these initial recordings, so these are assigned with low confidence.

In addition to the ipRGC types described above, ProA27 predominantly targeted RGCs which exhibit small somas and highly branched, compact dendritic arbors that stratify narrowly in the middle layer of the IPL (stratum 3), consistent with the class of ON-OFF small receptive field RGCs, which includes four subtypes. Based on rgctypes.org and MRCA, HD1/C13, HD2/C6 and UHD/C2 types are Tusc5+/FoxP2-(∼20% labeled cells, Figure 3C), and LED/C11 cells are Tusc5-/Foxp2-. Neuronal reconstructions most closely resembled the HD1 and HD2 subsets. These have very similar morphologies but can be separated physiologically based on their light responses (Jacoby and Schwartz 2017). We identified HD1 cells (n=2) characterized by a slow sustained spiking response to ON stimuli, and a delayed slow transient OFF response (Fig. 5A iv). The spatial tuning analysis revealed that ON spiking responses peaked about 200 µm and were greatly reduced by 400 µm indicating a small receptive field. A slight direction preference was seen in response to a drifting grating presented in 8 directions. We also found HD2 cells (n=2) characterized by a delayed spiking response to ON stimuli, and a delayed, but more sustained response to OFF. Similar to HD1, the spiking response during both the ON and OFF stimuli were greatly reduced at 400 µm indicating a small receptive field. HD2 cells also displayed aa slight direction preference.

**Figure 5.**
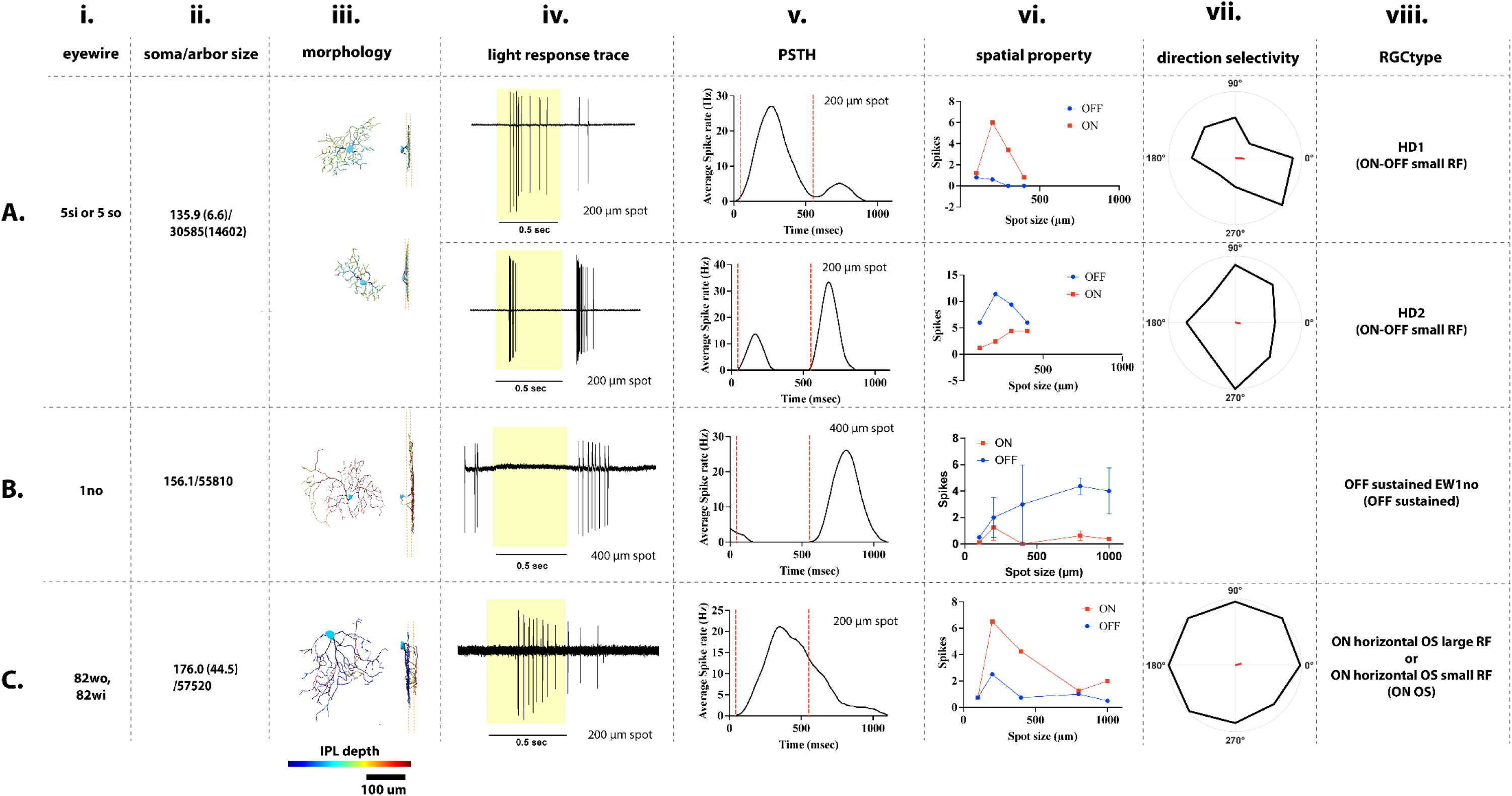
ProA27 labels RGCs with diverse morphologies and response properties. (Column i.) Putative cell types were identified based on morphology matching to the EyeWire database. (ii.) Soma and dendritic arbor areas were measured in projections from confocal stacks. (iii.) Examples of traced RGC’s matched to putative EyeWire types shown *en face* and from the side illustrate morphology and dendrite stratification. Patch clamp recordings from a separate set of retinas revealed light response properties including (iv.) spike traces, (v.) PSTH to a 1 Hz square-wave modulation with the indicated spot size, (vi.) average total spike counts to 1 Hz square-wave modulation during ON (red) and OFF (blue) plotted as a function of stimulus diameter, and (vii.) vector plot (black) showing total spike count for 8 different directions to a 1 Hz drifting grating with 200 µm bars. Small vector (red) indicated vector sum. These were used to match to (viii.) types in the RGCtype database. (A) Cells with small dendritic arbors of ∼30,000 μm^2^ stratified in layer 3 between the two cholinergic bands were difficult to distinguish morphologically between EyeWire type 5si or 5so. Light responses from patch clamp recordings suggested these included HD1 and HD2 cells based on spike traces with a 200 μm spot and temporal response. Note that HD1 had a higher average spike rate in the ON phase than OFF well HD2 cells had the converse higher OFF response than ON. (B) Cells matched with the 1no type stratified in the off layer with an average arbor size ∼55,000 μm^2^. These corresponded to cells from patch clamp recording with an OFF sustained response and were matched with the type OFF sustained EW1no. (C) ProA27 also labeled a bistratified RGC type with dendritic processes in layer 2 and 5. These were similar to EyeWire type 82wo or 82wi. In patch clamp recording, we identified a putative cell type with raw traces, preferred spot size, and temporal response consistent with ON horizontal OS large RF RGC type or ON horizontal OS small RF type.

**Figure 6.**
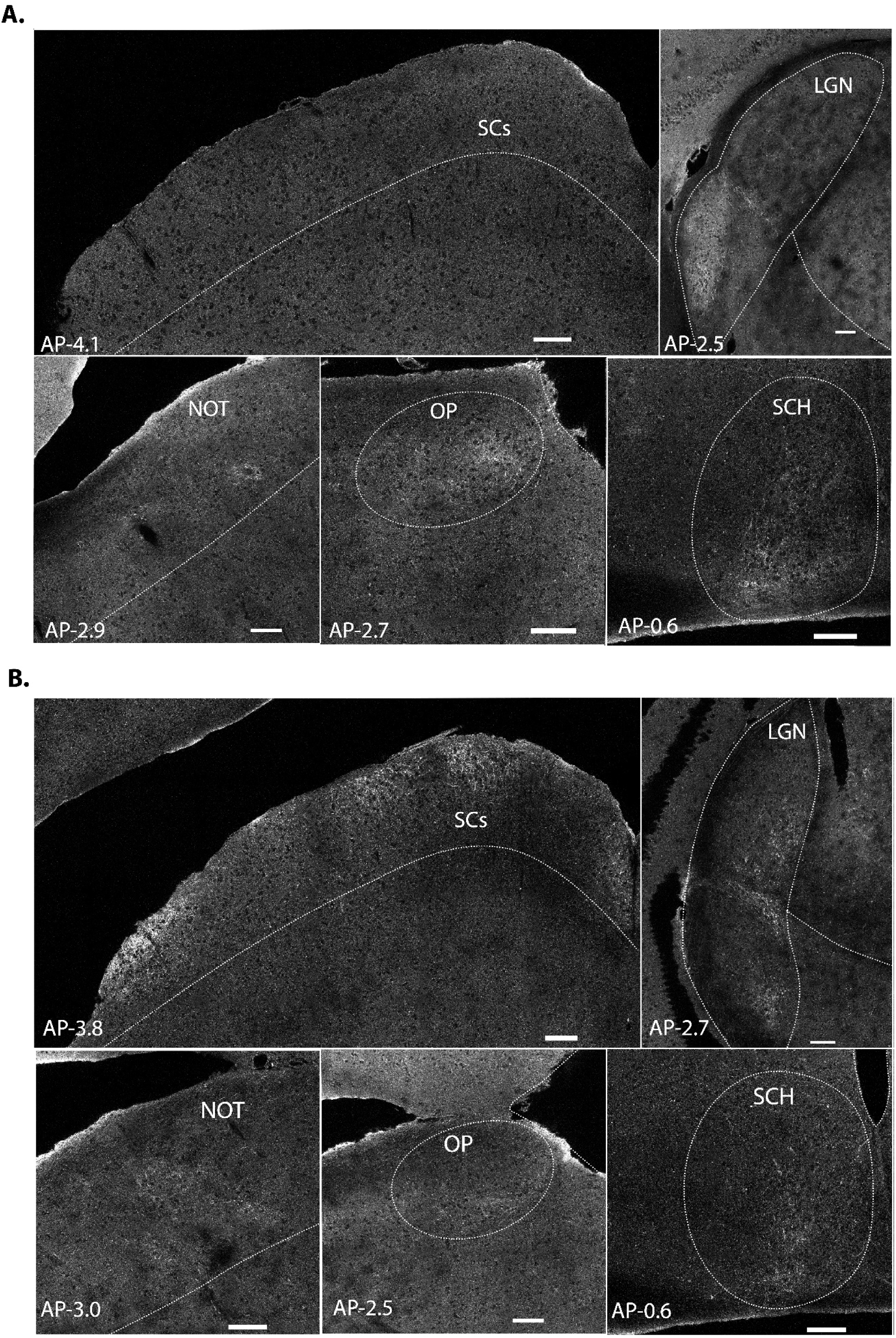
Distinct central projection patterns of ProA13- and ProA27-RGCs. (A) Coronal brain sections showing axonal projections from ProA13-RGCs visualized by native memGFP fluorescence. Labeled axons were observed predominantly in non-image-forming visual centers, including IGL, vLGN, the SCH, NOT, and OP. Notably, no signal was detected in SC, consistent with the non-image-forming identity of ProA13-targeted subtypes. (B) Brain sections from ProA27-AAVs injected mice reveal broader and more robust projection patterns. In addition to labeling non-image-forming centers (SCH, vLGN, OP, NOT), strong GFP signal was detected in image-forming regions such as dLGN and SC. Dashed lines delineate anatomical boundaries of the target regions. Anteroposterior (AP) coordinates from bregma are indicated in each panel. Scale bars, 50 μm.

Interestingly, the subtype with the highest endogenous expression of *Serpinb1b*, and thus the subtype predicted to be most consistently labeled with ProA27, was the LED/C11 type. These are morphologically similar to both HD and UHD types, but with a more uniform sustained response to both light onset and offset (Jacoby and Schwartz 2017). We did not detect cells that we could clearly match to this type.

Additional labeling in ProA27 was observed in RGCs, which have moderately sized somas and asymmetric dendritic fields stratifying in the deeper sublaminae of the IPL, which we matched to the EyeWire type 1no. Consistent with this, we found examples of OFF sustained cells (n=4) characterized by a slow sustained spiking response after stimulus offset. As very little baseline spiking response was observed, nothing was suppressed during ON stimuli. OFF spiking rates increased in response to larger spot sizes with very little suppression seen due to inhibition. A direction selectivity was not observed in these cells (Fig. 5B). Finally, we also recorded from several ON sustained cells in ProA27 (n=3). These were characterized by a slow onset and very sustained spiking response to ON stimuli that did not immediately respond to light offset. A direction selective preference was not seen in our cells. We tentatively assigned these to type ON horizontal OS large RF or EyeWire type 82wo, or type ON horizontal OS small RF or EyeWire type 82wi (Figure 5C).

### ProA13 RGCs primarily target non-image-forming pathways, whereas ProA27 RGCs project to both non-image-forming and image-forming visual pathways

RGCs are the sole output neurons of the retina, integrating visual input relayed from photoreceptors via interneurons and projecting their axons to several central targets. Among these, the LGN serves as a critical thalamic relay to the primary visual cortex and is essential for conscious visual perception (Monavarfeshani, Sabbagh, and Fox 2017; Seabrook et al. 2017). In mice, the LGN is anatomically subdivided into the dorsal LGN (dLGN), and ventral LGN (vLGN), separated by the intergeniculate leaflet (IGL), with each subregion receiving input from distinct RGC subtypes (Monavarfeshani et al. 2017). dLGN predominantly receives input from image-forming RGCs involved in spatial and contrast processing, whereas the IGL and vLGN innervated primarily by non-image-forming RGC types. The SCs also receives input from a range of RGC subtypes and contributes to both image-forming visual pathways, such as spatial localization and motion detection, and non-image-forming functions, including reflexive responses to ambient or salient visual stimuli (Basso and May 2017; Cang et al. 2018; Shang et al. 2015).

ProA13-RGCs mainly projected to the IGL, vLGN, SCH, nucleus of the optic tract (NOT), and olivary pretectal nucleus (OP), while notably absent from the SCs (Fig. 4A). The projection pattern aligns with known target preferences of the labeled subtypes. M1 ipRGCs target non-image-forming centers such as the SCH, vLGN, and OP, where they contribute to circadian entrainment and ambient light detection (Do 2019). Similarly, ON alpha RGCs have been reported to innervate both IGL and vLGN, forming part of thalamic relay circuits involved in brightness and motion encoding (Huang et al. 2019). PixON and ON vertical OS RGCs are typically enriched in the dLGN-projecting population (Kerschensteiner 2022), which may account for the low-level dLGN signal observed in our tracing experiments where they contribute to both image-forming and non-image-forming visual streams.

In contrast, the projection patterns of ProA27-RGCs reflect their broader subtype diversity. The presence of M1–M4 ipRGCs supports the strong labeling observed in SCH, vLGN, and OP, which are non-image-forming targets of melanopsin-expressing RGCs (Aranda and Schmidt 2021). W3 RGCs, known for their role in detecting local object motion, project robustly to the SCs (Kim et al. 2010; Zhang et al. 2012), matching the strong SCs signal seen in our dataset. Together, ProA27 labels a functionally diverse population of RGCs contributing to both image-forming and non-image-forming circuits, with prominent projections to the SCs and the dLGN. In conclusion, ProA13 and ProA27 drive expression in distinct RGC subtypes, with ProA13 preferentially targeting non-image-forming pathways projecting to IGL, vLGN, and SCH, while ProA27 labels a broader set of RGCs, including motion- and orientation-sensitive types, projecting prominently to the dLGN and the SCs.

## Discussion

In this study, we demonstrate that synthetic AAV promoters ProA13 and ProA27 enable selective labeling of molecularly and functionally distinct subsets of RGC types in the mouse retina. ProA13 predominantly labeled RGCs involved in non-image-forming visual circuits whereas, ProA27 targeted a broader array of RGC subtypes, and these cells projected extensively to both image-forming and non-image-forming brain regions. These findings highlight the utility of synthetic AAVs as flexible tools for accessing defined RGC population.

We employed a transcriptomic-guided strategy to choose synthetic AAV promoters using regulatory sequences derived from genes with enriched expression in RGCs (Jüttner et al. 2019). Single-cell RNA sequencing data indicated that *Nppb* is primarily expressed in RGCs, particularly in ipRGCs, ON alpha RGC, F-midi ON cells, and several novel transcriptomic clusters (Fig 1A). However, when used as a promoter in AAV (ProA13), the resulting *in vivo* expression pattern diverged from expectations.

Approximately half of GFP-positive cells were displaced amacrine cells (ACs), and horizontal cells (HCs) were also labeled. Among RGCs, ProA13 preferentially labeled M1 ipRGCs and ON-type RGCs such as ON alpha and PixON cells, whose dendrites stratify strongly in IPL sublamina 1, consistent with ON-layer targeting. A similar divergence between transcriptomic expression and AAV-mediated targeting was observed with *Serpinb1b* (ProA27). Although ProA27-RGCs expression was more restricted to RGC-specific. Instead of targeting Cluster 11 (C11) preferentially as suggested by gene expression, ProA27 labeled a diverse population of RGCs, including M1–M4 ipRGCs, a group of RGCs with dendrites stratified in layer three of the IPL, OFF sustained (EW1no), and ON horizontal OS RGCs. Approximately 73% of ProA27-RGCs were labeled in our panel of molecular markers (Fig. 3E), while the remaining population likely includes novel or transitional subtypes. A notable subset of ProA27-labeled RGCs exhibited distinct dendritic stratification in IPL layer 3, a lamina associated with motion-sensitive RGCs, including UHD, HD1, HD2, and LED cells (Jacoby and Schwartz 2017). Our patch clamp recording from these cells were most consistent with HD1 and HD2 subtypes. Indeed, we found that some of RGCs stratified in the IPL 3 were Tusc5-positive. However, we also observed Tusc5-negative cells stratifying in the same layer, which could correspond to LED/C11 cells, although we did not clearly capture these in our patch clamp recordings. These findings underscore both the strength and limitations of using transcriptome-informed promoters for cell-type-specific AAV targeting. While synthetic promoters based on single-gene expression profiles can enrich for particular RGC subtypes, actual expression patterns in vivo may diverge due to cell-type-specific differences in promoter accessibility, chromatin context, or AAV serotype tropism. Despite these challenges, ProA13 and ProA27 provide effective and reproducible genetic access to distinct RGC populations, enabling studies of their structural and functional properties.

By combining these synthetic promoters with membrane-targeted GFP and confocal imaging, we were able to visualize not only RGC morphology but also central axonal projections. This AAV-based approach offers advantages over transgenic mouse lines, including simplified implementation, flexible design, and compatibility with combinatorial strategies. Although each promoter labeled multiple RGC subtypes, potentially limiting subtype-specific resolution, we observed differences in projection patterns between ProA13- and ProA27-RGCs. ProA13-RGCs projected primarily to non-image-forming centers such as IGL, vLGN, SCH, OP, and NOT, with negligible input to sSC (Fig. 4A). In contrast, ProA27-RGCs projected broadly to both image-forming targets (dLGN, sSC) and non-image-forming structures (vLGN, SCH, OP) (Fig. 4B). A particularly notable difference was observed in the sSC. The sSC, a laminated structure in the midbrain, integrates multimodal sensory inputs. We found dense ProA27-labeled projections to the superficial layers of the sSC, consistent with previous reports showing that RGCs responsive to local object motion, such as W3 RGCs (Jacoby and Schwartz 2017; Kim et al. 2010), preferentially innervate the sSC. In contrast, alpha RGCs and ipRGCs tend to target deeper SC layers (Dhande and Huberman 2014), aligning with the sparse sSC labeling in deeper SCs layer.

The ability to selectively label and access distinct RGC subtypes using synthetic AAV promoters such as ProA13 and ProA27 opens new opportunities for targeted interrogation of visual circuits. These tools facilitate cell-type-specific access without the need for complex transgenic models, enabling rapid deployment across diverse experimental contexts. The differential projection patterns observed between ProA13 and ProA27 further allow for the dissection of parallel visual processing pathways and their contributions to image-forming versus non-image-forming functions. Moreover, this AAV-based approach can be extended to deliver optogenetic actuators, calcium indicators, or gene-editing tools, providing a versatile platform for functional circuit manipulation. Altogether, this strategy significantly enhances our ability to probe how specific RGC types contribute to vision and brain-wide sensory processing.

## Methods and Materials

### Animals

All mice were maintained on C57BL/6 background. Adult mice of both sexes aged postnatal days (P) over 42 were used in this study. Animals were provided with food and water ad libitum and maintained on a 12-hour light/12-hour dark cycle. All procedures were approved by the Wayne State University Institutional Animal Care and Use Committee (IACUC).

### AAV Preparation

AAV vectors driven by synthetic promoters A13 (*Nppb*) and A27 (*Serpinb1b*) were constructed starting with plasmids described by Jüttner et al. 2019, pAAV-ProA27-CatCh-GFP-WPRE (Addgene plasmid # 125910; http://n2t.net/addgene:125910; RRID:Addgene_125910) and pAAV-ProA13-CatCh-GFP-WPRE (Addgene plasmid # 125896; http://n2t.net/addgene:125896; RRID:Addgene_125896) were gifts from Botond Roska. Each construct was modified by subcloning cDNA for a membrane-targeted GFP (memGFP) reporter in place of CatCh-GFP. To produce viral particles, 293AAV cells were cultured in DMEM supplemented with 10% heat-inactivated fetal bovine serum (FBS), 1% penicillin-streptomycin, and 2 mM L-glutamine. Cells were transfected at 70–80% confluency with the transfer plasmid, pAAV-AAV2 capsid plasmid, and pHelper plasmid using a calcium phosphate method in IMDM.

Viral particles were harvested 72 hours post-transfection, treated with sodium deoxycholate and Benzonase nuclease, and purified using heparin column chromatography with stepwise NaCl elution. Concentrated viral preparations were filtered through 0.22 μm filters and stored at –80°C until use.

### Intraocular injections

Adult mice were anesthetized with 3–5% isoflurane in oxygen and a small puncture was made at the peripheral sclera using a 27-gauge needle to create an entry site, and a microliter syringe (10 μL, PN:7635-01, Hamilton company) connected to a blunt needle (33-gauge, PN:7762-06 Hamilton company) was inserted through the sclera into the vitreous chamber. A total volume of 1.5 µL of AAV solution was injected per eye. After injection, the micropipette was carefully withdrawn, and Proparacaine hydrochloride ophthalmic solution (NDC 24208-730-06) was applied to the eye. Mice were monitored during recovery and returned to their home cages once fully ambulatory. All procedures were conducted in accordance with institutional animal care and use guidelines and approved animal protocols.

### Retinal and Brain Preparation and Immunohistochemistry

Mice of either sex were sacrificed via CO₂ asphyxiation and transcardially perfused with PBS, followed by 4% paraformaldehyde (PFA).

#### Whole-Mount Retina Preparation

Eyes were enucleated and dissected to remove the cornea and iris. Retinas were fixed in 4% PFA for 4–8 hours at 4°C, then carefully isolated and stored in 1X phosphate-buffered saline (PBS) at 4°C until staining. For primary antibody staining, retinas were washed once or twice in 1X PBS to remove debris and incubated in 250 μL of primary antibody diluted in blocking solution (2.5% bovine serum albumin [BSA], 0.25% Triton X-100 in PBS) for 24–72 hours at 4°C, depending on the antibody. Following incubation, retinas were washed three times in PBS (30 minutes each) to remove unbound primary antibodies. Secondary antibodies, diluted in the same blocking solution, were then applied for 24 hours at 4°C. Retinas were washed three additional times in PBS (30 minutes each) and stored in PBS at 4°C until mounting. For imaging, retinas were mounted on microscope slides using 80% glycerol. After imaging, coverslips were carefully removed, and retinas were returned to PBS for storage or further processing.

#### Cryosection Immunostaining

Fixed retinas were cryoprotected in 30% sucrose, embedded in Tissue-Tek OCT compound (Sakura), and sectioned at 12 μm thickness onto Superfrost Plus microscope slides. Slides were incubated in blocking solution (2.5% BSA and 0.25% Triton X-100 in PBS) for 1–2 hours at room temperature. After blotting off the blocking solution, primary antibodies diluted in blocking buffer were applied directly onto the sections. Slides were covered with trimmed parafilm and incubated overnight at 4°C.

The following day, slides were washed three times in 1X PBS (∼5 minutes each) and incubated with secondary antibodies diluted in PBS for 1 hour at room temperature. Slides were washed again three times in PBS, with DAPI (1 μL per 1X PBS) added during the final rinse. After a brief rinse in distilled water, slides were allowed to dry, and ∼60 μL of mounting medium was applied before sealing with coverslips for imaging.

#### Antibodies

The following antibodies were used for immunohistochemistry:

| **Antigen** | **Host Species** | **Dilution** | **Source** | **Catalog Number** | **RRID** |
| --- | --- | --- | --- | --- | --- |
| RBPMS | Guinea Pig | 1:1000 | Millipore | ABN1376 | AB_2687403 |
| Pax6 | Mouse (IgG1) | 1:100 | DSHB | pax6 | AB_528427 |
| VACHT | Rabbit | 1:200 | Abcam | ab235201 | AB_3697034 |
| ChAT | Goat | 1:500 | Sigma-Aldrich | AB144P | AB_2079751 |
| Parvalbumin (PV) | Rabbit | 1:500 | Swant | PV27 | AB_2631173 |
| eGFP | Chicken | 1:2000 | Aves Labs | GFP-1010 | AB_2079751 |
| Melanopsin | Rabbit | 1:5000 | Advanced Targeting Systems | UF006 | AB_2314781 |
| SMI32 | Mouse (IgG1) | 1:1000 | Covance | SMI-32R-100 | AB_509997 |
| CART | Rabbit | 1:2000 | Phoenix Pharmaceuticals | H-003-62 | AB_2313614 |
| Tusc5 | Mouse (IgG1) | 1:500 | Santa Cruz Biotech | Sc-377025 | AB_3697030 |
| FoxP2 | Sheep | 1:1000 | R&D Systems | AF5647 | AB_2107133 |

All secondary antibodies were species-specific and conjugated to Alexa Fluor dyes (Alexa Fluor 488, 568, or 647), and were used at a dilution of 1:500 in blocking solution.

#### Brain Sectioning for Imaging

After fixation for overnight at 4°C, brains were sectioned coronally at 100 μm thickness using a vibratome (e.g., Leica VT1000S). Free-floating sections were stored in 1X PBS at 4°C until immunostaining. The same antibody labeling and washing protocol used for retina cryosections was applied for brain sections, with minor adjustments in volume and incubation time as needed.

### Imaging and morphological reconstruction

Following immunostaining, samples were imaged using either an epifluorescence microscope (Leica) or a confocal laser-scanning microscope (Leica), depending on the experimental requirements. High-resolution z-stack images of individual RGCs were acquired using 40× water-immersion objectives. For confocal imaging, optical sections were captured at step sizes of 0.4–0.5 μm to ensure accurate three-dimensional reconstruction. RGC morphology was reconstructed from confocal image stacks using Imaris software (Bitplane). Membrane-tagged GFP signals were rendered in 3D using the “Filament Tracer” module. VAChaT or ChAT immunostaining was used as a reference to align dendrites within the (IPL. Quantitative and qualitative morphological comparisons were made across experimental groups.

Dendritic arbor size was calculated by outlining the outermost boundaries of the dendritic field using a polygonal selection tool in Fiji (ImageJ), and the enclosed area was measured. Brain sections containing axonal projections were imaged using a confocal laser-scanning microscope. Sections were mounted in mounting medium without additional staining, as native GFP fluorescence from membrane-tagged reporters was sufficient for visualization.

### Electrophysiological measurement

#### Retinal preparation

The experimental techniques used were described in detail in previous studies (Bohl et al. 2022; Ichinose, Fyk-Kolodziej, and Cohn 2014). In brief, both male and female adult C57BL/6 background were used.

Prior to tissue collection, all animals were kept in darkness overnight. They were then euthanized using carbon dioxide followed by cervical dislocation, after which their eyes were enucleated. Retinas were carefully dissected under a microscope under dark-adapted conditions using infrared visual aids. The tissue preparation was carried out in a HEPES-buffered solution containing 115 mM NaCl, 2.5 mM KCl, 2.5 mM CaCl₂, 1.0 mM MgCl₂, 10 mM HEPES, and 28 mM glucose, with the pH adjusted to 7.37 using NaOH. Throughout the dissection, a chilled and continuously oxygenated solution was used. Retinal samples were then stored in an oxygenated dark box at room temperature.

#### Whole-cell recordings

Whole-cell patch-clamp recordings were conducted on ganglion cell somas expressing Green Fluorescent Protein (GFP) in wholemount retinal preparations. A two-photon microscope (Borghuis Instrument, Louisville, KY) equipped with a laser system (Axon 920, Coherent, Inc, CA) with a Scan Image 3.8 software or fluorescence microscopy (Scientifica and CoolLED, UK) was used for GFP visualization.

Retinal tissue was secured using a platinum horseshoe-shaped frame and nylon threads. Light-evoked spikes were recorded at the cells’ resting membrane potential, while light-evoked excitatory postsynaptic currents (L-EPSCs) were measured at a holding potential of −55 mV (at ECl) during voltage-clamp recordings. All recordings were carried out at a temperature range of 32–34°C in Ames’ medium buffered with sodium bicarbonate (NaHCO₃, Millipore-Sigma, St. Louis, MO, USA) and continuously oxygenated with a 95% O₂ / 5% CO₂ gas mixture to maintain a physiological pH of 7.4. Patch pipettes were fabricated from borosilicate glass (1B150F-4; WPI, FL, USA) using a P1000 micropipette puller (Sutter Instruments, CA, USA), yielding resistances between 4 and 9 MΩ. Data acquisition and stimulus generation were managed using Clampex software (pClamp 11) and a MultiClamp 700B amplifier (Molecular Devices, CA, USA). Signals were digitized via an Axon Digidata 1550B interface (Molecular Devices), low-pass filtered at 1 kHz with the MultiClamp’s four-pole Bessel filter and sampled at rates ranging from 2 to 5 kHz.

#### Light stimulation

Retinal wholemount tissues were light-adapted at 1 × 10^5^ photons/μm2/s in the recording chamber. Recorded ganglion cells were presented with a light from the bottom of the microscope stage through a condenser (500nm, 1 × 10^7^ photons/μm^2^/s, 150μm diameter). A series of light stimuli is as follows: 1. A step light for 500 ms from the background light to dark (1 × 10^4^ photons/μm^2^/s) and bright (1 × 10^6^ photons/μm^2^/s). 2. White circular pattern from the diameters of 100 µm to 1000 µm. 3. Eight directional grating (grating speed: 1Hz; bar size: 3636 µm).

#### Data analysis

The number of spikes, spike frequency (Hz), and the amplitude of L-EPSCs were measured using the Clampfit (Molecular Devices) and custom-coded MatLab (Mathworks).

